# The interplay of trophic interactions and game dynamics gives rise to life-history trade-offs, consistent personalities, and predator-prey and aggression power laws

**DOI:** 10.1101/2024.02.13.580043

**Authors:** Mohammad Salahshour

## Abstract

By introducing a simple model of ecological interactions where individuals interact trophically, and through a game dynamic, I show that the dissipative flow of resources can derive evolution and lead to the emergence of a scale-invariant ecosystem exhibiting a wide range of mean and fluctuation scaling laws that govern trophic interactions and game dynamics. The eco-evolutionary approach suggests life history trade-offs are a natural consequence of ecological dynamics and, combined with the non-equilibrium dynamic, lead to the evolution of consistent personalities. Aggressiveness and personality consistency depend on trophic position, and predators evolve a higher aggressiveness and starker personality differences.

**Author summary:** Throughout the history of life, the flow of energy across ecosystems has contributed to the evolution of complex forms of life and strikingly universal patterns on a large scale. However, it is not clear what factors lead to such universal patterns and how they relate to evolution. Simple mathematical models suggest that the dissipative flow of resources through ecosystems leads to self-organization in a critical state with scale-invariant avalanches of activity. The scale-invariant structure of ecosystems results in a complex set of scaling laws governing the structure and dynamics of populations. The same non-equilibrium ecological dynamics derive evolution and account for the evolution of individuals’ behavioral differences and consistent personalities.

## Introduction

Ecological dynamics and evolution are derived from the constant flow of resources into the ecosystems and their dissipation along the food web gradient. In physical systems, such a dissipative flow of energy can lead to the system’s self-organization into a critical state with scale-invariant avalanches of activity, self-similar and fractal structures, and power law distributions [1–10]. Indeed, hallmarks of criticality, such as fractal structures [11], power law relations of predator and prey biomass [12], scaling in food webs [13, 14], power law biomass distribution on earth [15–17], scaling of metabolic functions [18–20], species-area relationships [21], or Taylor power laws governing the mean and fluctuations of populations [22–24], to name a few, have been observed in ecological systems. However, it is not clear how these scale-invariant patterns relate to the flow of energy in ecosystems, and what similarities they may have with self-organized criticality.

Recently, it has been shown that such a dissipative flow of energy can have profound consequences and explain some of the most striking large-scale ecological patterns shown by diverse ecosystems across the globe [25]. A simple agent-based model of predator-prey interactions where prey consume primary resources and predators feed on prey, suggests the scale-invariant spatial distribution of predators and prey, as well as a diverse range of fluctuation-scaling laws and predator-prey power laws, can be a simple consequence of the flow of energy across ecosystems. However, while such patterns can emerge in a purely ecological model of trophic interactions, this observation can have profound consequences for evolutionary dynamics as well. The accumulating evidence of rapid evolution and widespread feedback between ecological and evolutionary dynamics has given rise to the view that evolution can naturally result from ecological dynamics, and in principle, at the time scales comparable to the time scale of ecological changes [26–30]. This body of evidence calls for such a theoretical shift in evolutionary theory, which is suggested to usher in a new synthesis of ecology and evolution [30, 31], with possibly a high impact on our understanding of ecological and evolutionary dynamics as a self-organizing complex system [32–35]. A fruitful mathematical approach to investigating the interplay between ecological dynamics and evolution is recently introduced in the context of evolutionary game dynamics, where it is shown that evolution can result naturally out of the flow of energy in ecosystems and organisms’ growth, and can be studied without appealing to fitness-based selection [36].

Here, by combining game dynamics within species and trophic interactions between species, I address the evolution of within-species interactions as a result of ecological dynamics and reveal how large-scale eco-evolutionary patterns, life history trade-offs, and consistent personalities can arise as a result of eco-evolutionary dynamics. To do so, I consider two populations, prey, and predators, interacting with each other through trophic interactions and (possibly) within populations by playing *n_g_* games. Individuals accumulate resources due to interactions, which they use to pay for their metabolic costs, grow, and reproduce. Individuals die when they go out of resources, reproduce when they reach enough resources for reproduction, and inherit their strategies in the game they play, subject to mutations. When individuals play no game, *n_g_* = 0, the model reduces to a purely ecological model of trophic interactions. The study of the purely ecological dynamics of prey and predator interactions shows that the flow of resources in trophic systems and its dissipation along the food web gradient gives rise to a scale-invariant distribution of individuals with a scale-invariant avalanche of activities resembling those observed in self-organized critical systems. A consequence of the scale-invariance of trophic systems is the scale-invariant spatial and temporal distribution of prey, predators, and basal resources, which leads to a wide range of scaling laws. While the recently discovered predator-prey power law [12] and the famous Taylor’s power law [22] are among the scaling laws predicted by the model, the model suggests both these laws can be extended to a large set of predator-prey mean and fluctuation scaling laws, according to which, mean and fluctuations of several variables, pertaining to the same or different populations scale with each other.

Once individuals inherit traits that affect their behavior, i.e., *n_g_ >* 0, evolution naturally results from ecological dynamics and inherits its scale-invariant structure. However, in contrast to standard frameworks of evolution, such as those used to study game dynamics (e.g., [37–39]), and consistently with a new eco-evolutionary approach [36], no fitness function is explicitly defined. Instead, evolution takes place as a direct consequence of ecological interactions and is inseparable from the latter. I consider a case where individuals play a modified version of the Hawk-Dove game where they can be aggressive or non-aggressive. Eco-evolutionary dynamics leads to the evolution of life history trade-offs: aggressive individuals live shorter lives and reproduce more, while nonaggressive ones live longer lives and reproduce less. I show that such life-history trade-offs can lead to the evolution of animal personalities. While this finding is consistent with past findings [40, 41] and the widespread rationale [42], in contrast to past works, life-history trade-offs are not introduced as a model assumption but evolve naturally as a result of eco-evolutionary dynamics. Furthermore, the model suggests that, while necessary, life history trade-offs are insufficient for the evolution of consistent personalities. Rather, the evolution of personalities crucially depends on the nonequilibrium dynamics, which derives the system out of the physical and game (Nash) equilibrium. The model predicts that predators evolve to be more aggressive than prey and show stronger personality consistency. Furthermore, aggressiveness endows indirect predatory or antipredatory benefits, which can favor its evolution in both prey and predators. These indirect benefits weaken the prey’s life-history trade-offs, which explains why prey show a weaker personality consistency. Finally, population viscosity hinders the evolution of consistent personalities by suppressing the evolution of aggressiveness due to its positive role in promoting altruistic behavior [43, 44].

Importantly, the model predicts a new class of scaling laws which can be called aggression scaling laws, according to which the number of deaths due to aggressive encounters, as well as densities and birth rates of individuals with different aggressiveness, show a rich set of mean and fluctuation scaling laws. By studying different variants of the model, where reproduction occurs through mitosis or when offspring acquire only a small fraction of its parents’ resources, and studying different parameter values and motility patterns, I show that self-organized criticality and scaling laws are highly robust features of the eco-evolutionary dynamics. However, the precise value of the scaling exponents may depend on different factors. While aggression power laws are yet to be empirically discovered, the recent discovery of a sublinear predator-prey power law [12] motivates a study of the factors affecting the predator-prey power law exponent. I show that small offspring size, predator competition, and surprisingly, prey competition, low predator conversion efficiency, small body size, and high prey relative motility all can contribute to a sublinear predator-prey power law and lead to a bottom-heavy food web with prey favoring density-dependency of life histories, according to which preys live longer lives and reproduce less in higher densities. Finally, the model sheds light on the life histories of the individuals by showing that small body size, competition, and smaller offspring size, lead to faster life history patterns in higher densities in both prey and predators. While the density dependence of life histories has been subject to some research [45–47], these findings suggest scaling laws may be at work in the density dependence of life histories and relate to the food web control and between populations and within populations scaling laws.

### The model

We consider an eco-evolutionary model of trophic interactions combined with a game dynamic (Fig. 1). Preys and predators move on a grid of linear size *L*. Each agent *α*, prey or predator has an internal resource, *ϵ_α_*, which increases by gaining resources, *κ_α_*, and decreases by a constant rate, *η*, due to paying for metabolic costs. Thus, the internal resource of agent *α* at time *t* evolves according to:

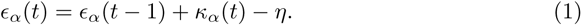

**Fig 1.**
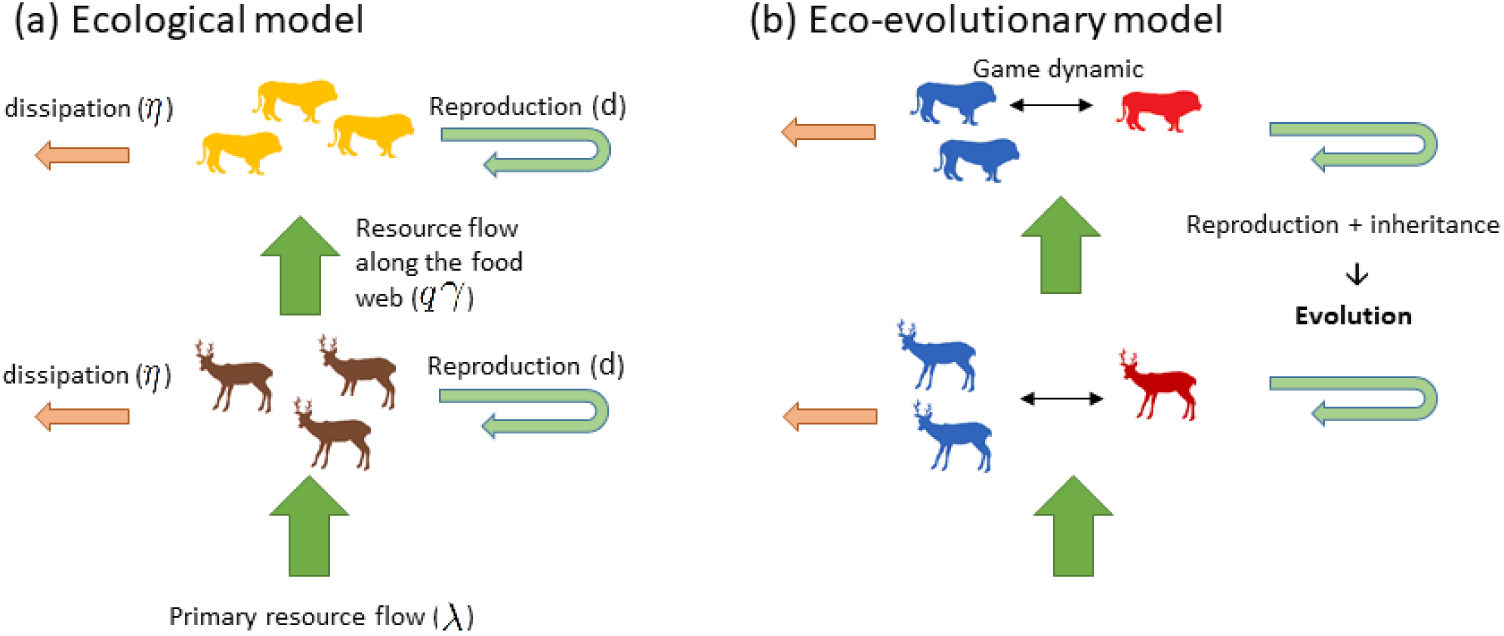
Illustration of the models. (a): In the purely ecological model, preys consume primary resources injected into the system with a rate *λ*, and predators convert prey into energy with a conversion efficiency, *γ*. Individuals’ energy reduces by a constant consumption rate, *η*, and they reproduce when their energy reaches a threshold, *d*. (b) By allowing the inheritance of strategies, ecological dynamics lead to evolution. In the eco-evolutionary model considered here, in addition to trophic interactions, individuals play a (Hawk-Dove) game with their conspecifics and inherit their strategy subject to mutations.

Preys acquire resources by consuming primary resources, which are regenerated on each site ***r***, with a rate *λ*(***r***). Basal resources available on a site are divided equally among all the prey visiting a site, subject to the condition that a prey can not acquire more resources than its maximum size *d*. Predators capture preys to gain resources. When a predator visits a site where prey lives, the predator captures a prey with probability *q*, and its internal resource increases by an amount *γϵ_α_*, where *ϵ_α_* is the internal resource of its prey, and *γ*is the predator’s energy conversion efficiency [48, 49] or more precisely, specific growth [50]. Preys and predators die if their internal resource reaches a subsistence level, Δ, and reproduce if their internal resource reaches the threshold *d*. For reproduction, we study cases where the parent’s resource is equally divided between the individual and its offspring (mitosis) and cases where an amount *b_o_* is transferred to the offspring.

In the evolutionary version of the model, in addition to the trophic interaction, individuals play *n_g_* games with others in the same population. For this purpose, at each time step, all the individuals on a lattice site are paired to play a modified version of the hawk-dove game. In this game, individuals have one of two strategies, hawk (aggressive) and dove (nonaggressive). When a hawk meets a dove, the hawk steals an amount of resource *r* from the dove. When two hawks or two doves meet, no resource is transferred. However, when two hawks meet, each individual has a *d_t_* probability of death due to a confrontation. Upon reproduction, individuals inherit the strategies of their parent subject to mutations with a probability *ν*.

When individuals play no game, *n_g_* = 0, the model reduces to a purely ecological model of trophic interaction (Fig. 1(a)). When *n_g_ >* 0, the model becomes an eco-evolutionary model where individuals’ strategies in the game are subject to evolution (Fig. 1(b)). However, in contrast to standard models of evolution, evolution is a natural consequence of ecological interactions and individuals’ reproduction, and no fitness function is explicitly defined. We study both the purely ecological version of the model and the eco-evolutionary version. For the sake of simplicity, we do most of the analysis using a simple version of the model by setting Δ = 0 and with mitosis. However, all the results are strikingly robust for different model modifications.

## Results

### Scale-invariance of eco-evolutionary dynamics and scaling laws

In both the ecological and eco-evolutionary models, as the resource regeneration rate increases, the system shows successive phase transitions from a phase where predators do not survive to a phase where prey and predators co-exist in equilibrium, and a phase where the system exhibits non-equilibrium dynamics with traveling waves of prey and predators (See the Supplementary Information, SI. 3). Furthermore, the model exhibits the paradox of enrichment [51–53]: for too high resource regeneration rates, the dynamics become unstable in a homogeneous environment. However, resource heterogeneity restores the system’s stability [25] (see SI. 3). Thus, here, we consider a heterogeneous environment with a linear distribution of resources in which the resource regeneration rate *λ*(***r***) on a site ***r*** = (*x, y*) is given by *λ*(***r***) = 2*λx/L*.

A striking feature of the non-equilibrium dynamics is scale-invariant avalanches of activity and power-law spatial distribution of the individuals. These features hold in the purely ecological model and are further duplicated when evolution takes place. The power-law distribution of avalanches of activity in the system for *n_g_* = 1 is studied in Fig. 2. All the results hold in the purely ecological model (*n_g_* = 0) and different variations of the model and parameter values (see the Supplementary Information, SI. 4). Fig. 2(a), from top to bottom, shows the amount of resources, *n_r_*, the number of prey, *n_p_*, and the number of predators, *n_π_* on a lattice site. As a result of the individuals’ interactions, all these variables show fluctuations. These fluctuations are reminiscent of avalanches of activity in self-organized critical systems [1–3]. To assess whether this similarity is profound, we define avalanche time as the time duration of an avalanche and avalanche size for resources, prey, and predators, respectively, as the total amount of resources, the number of prey or predators who have lived during an avalanche (see SI. 2). These quantities are plotted in Figs. 2(b) and 2(c) for different values of resource regeneration rate and show a scale-invariant frequency-size distribution. Postulating a universal scaling form, *λ^−β_x_^ f_λ_*(*xλ^−α_x_^*), where *α_x_* and *β_x_* are proper exponents, and *x* refers to resource, *r*, prey, *p*, and predator, *π*, avalanche time or size, the data for all the resource regeneration rates collapse into a universal scaling function.

**Fig 2.**
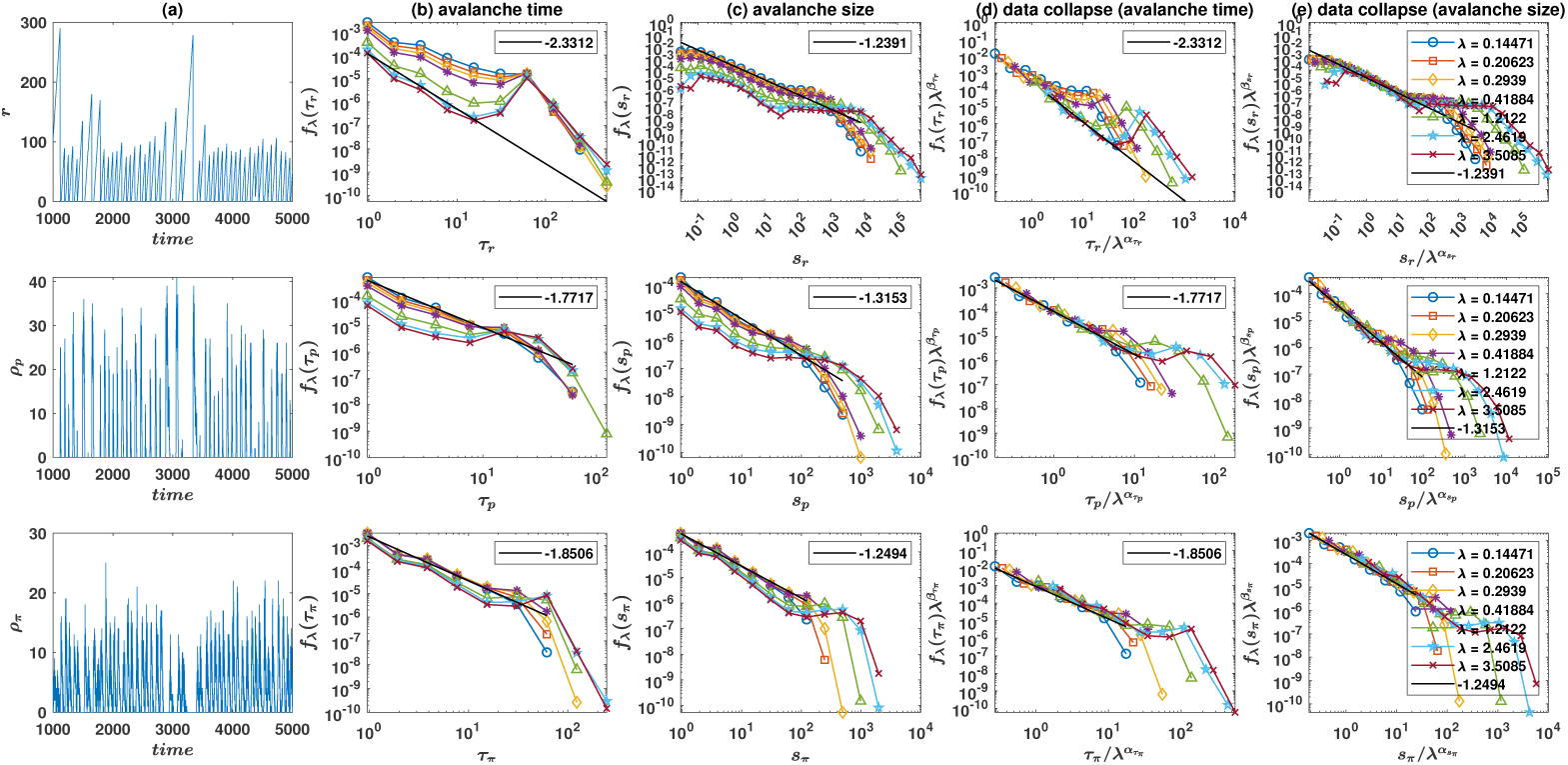
Power-law avalanches of activity in eco-evolutionary dynamics. (a): The amount of resources, *r*, (top), the number of prey, *ρ_p_*, (middle), and predators, *ρ_π_* (bottom) on a lattice site. (b): An avalanche for, respectively, resources, preys, and predators is defined as an event where a site remains active (i.e., contains nonzero, respectively, resource, prey, or predators). Avalanche time is the time duration of an avalanche, and avalanche size is the total amount of resources or the number of prey or predators who have visited a site during an avalanche. Both avalanche time, *τ*, (b) and size, *s*, (c), for different values of the resource regeneration rates, show a power law distribution. (d) and (e): When the rescaled avalanche time and size distribution *f_λ_*(*τ_x_*)*λ^β_τ_x__^* and *f_λ_*(*s_x_*)*λ^β_s_x__^*, are plotted as a function of, respectively, rescaled avalanche time *τ_x_/λ^α_τ_x__^* and size *s_x_/λ^α_s_x__^*, the data for different values of the resource regeneration rate collapse into a single universal curve. exponent values used for data collapse: *α_τr_* = −0.85, *β_τr_* = −0.9, *α_τp_* = −0.85, *β_τp_* = −0.71, *α_τπ_* = −0.65, *β_τπ_* = −0.71, *α_sr_* = −0.42, *β_sr_* = 0.78, *α_sp_* = −0.85, *β_sp_* = −0.4, *α_sπ_* = −0.85, *β_sπ_* = −0.71. Parameter values: *η* = 0.05, *γ* = 0.5, *d* = 2, *q* = 0.5. An eco-evolutionary model with *n_g_* = 1, *r* = 0.4, *d_t_* = 0.05, and *ν* = 10*^−^*^3^ is considered. In (a), *λ* = 0.8507.

Another manifestation of the scale-invariance of the eco-evolutionary dynamics is the scale-invariant distribution of the amount of resources, the number of prey, and predators on a lattice site. These distributions for different values of the resource regeneration rate are plotted in Fig. 3 and show a power law distribution for small sizes with a sharp cut-off at large sizes. In this case, too, the distribution resulting from different resource regeneration rates collapses into a single universal curve upon proper rescaling. This can be seen in Fig. 3(a) to (c), where the data collapse resulting from plotting the rescaled frequency *f_λ_*(*n_x_*)*λ^β_x_^* as a function of the rescaled size, *n_x_/λ^α_x_^* is shown. Furthermore, the cut-off, which is the maximum number of individuals observed on a lattice site in the system, shows scaling with *λ*, with a decreasing exponent going to the upper levels of the food web.

**Fig 3.**
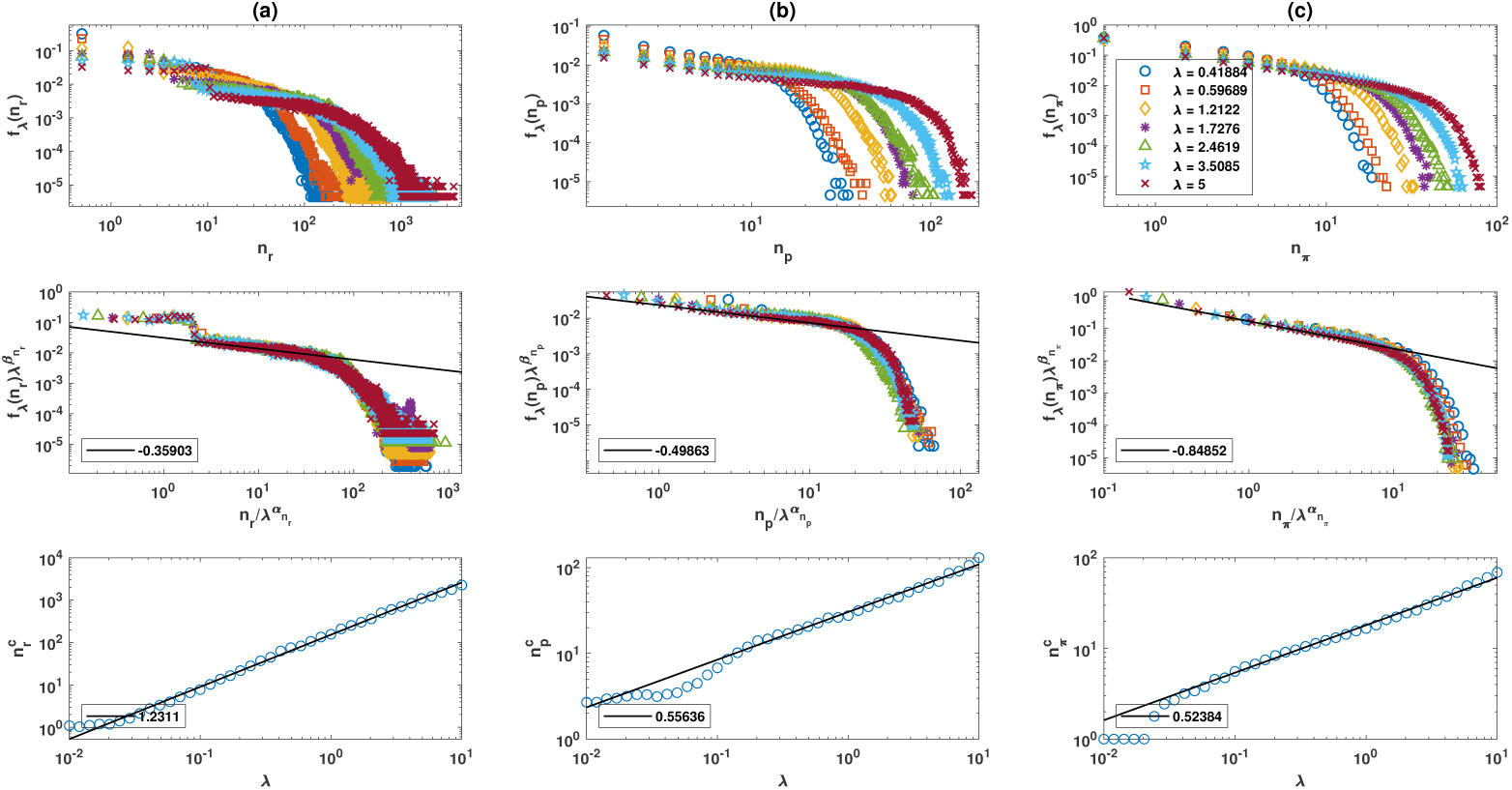
Scale-invariance of the spatiotemporal distribution of basal resources, prey, and predators. The distribution of the amount of resources (a), the number of prey (b), and predators (c) on a lattice site for different values of the resource regeneration rate, *λ* (top row). All the distributions show a scaling regime in small sizes with a sharp cut-off for large sizes. Plotting the rescaled distribution, *f_λ_*(*n_x_*)*λ^β_x_^* as a function of the rescaled size, *n_x_/λ^α_x_^* the distributions for different values of *λ* collapse into a single universal curve (middle row). The cut-off shows scaling with the resource regeneration rate (bottom row). Exponent values used for the data collapse: *α_r_* = 1, *β_r_* = 1, *α_p_* = 0.75, *β_p_* = 0.65 *α_π_* = 0.75, *β_π_* = 0.8. Parameter values: *η* = 0.05, *γ* = 0.5, *d* = 2, *q* = 0.5. An ecoevolutionary model with *n_g_* = 1, *r* = 0.4, *d_t_* = 0.05, and *ν* = 10*^−^*^3^ is considered.

The data collapse of the spatiotemporal distributions has profound consequences for ecological dynamics by giving rise to a rich set of scaling laws [25]. As detailed in the Methods, the universal scaling form of the spatial distribution of a generic variable *v* and the scaling of the cut-off with *λ* predicts *v* shows a power law relation with *λ*. This gives rise to a large set of primary scaling relations, some of which are plotted in Fig. 4, top rows, (see the Supplementary Information, SI. 8 and SI. 9 for a more detailed list). The density of prey *ρ_p_* and predators, *ρ_π_*, and their internal energy, *M_p_* and *M_π_*, prey and predator per-area birth rate, *B_p_* and *B_π_*, prey production defined as the number of prey caught by predators, *P_p_*, the per-area death rate of prey and predators in aggressive encounters *D_p_* and *D_π_*, the density of prey and predators with dove (*ρ_p_*_(*d*)_ and *ρ_π_*_(*d*)_) and hawk, (*ρ_p_*_(_*_h_*_)_ and *ρ_π_*_(_*_h_*_)_) strategy, and their per-area birth rate, (*B_p_*_(*d*)_, *B_π_*_(*d*)_, *B_p_*_(_*_h_*_)_, and *B_π_*_(_*_h_*_)_) show scaling with the resource regeneration rate, *λ*. Furthermore, the temporal (and spatial) fluctuation of the same quantities show scaling with *λ* (Fig. 4, bottom rows).

**Fig 4.**
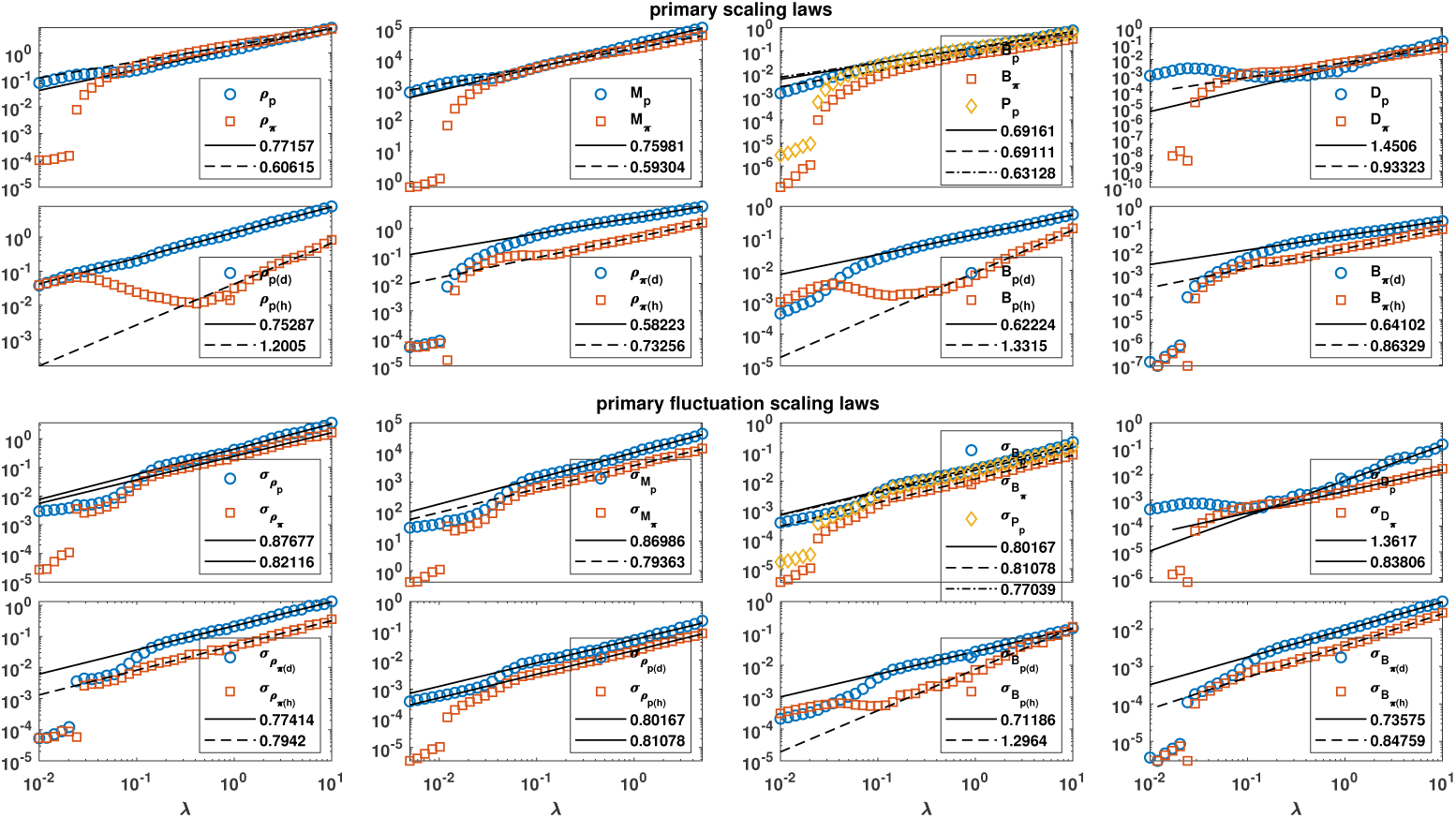
Primary scaling laws. The eco-evolutionary dynamics exhibits primary scaling laws according to which mean and fluctuations of several variables show scaling with the resource regeneration rate. Prey and predator densities, *ρ_p_*, and *ρ_π_*, total internal energy *M_p_*, and *M_π_*, per-area birth rate *B_p_* and *B_π_*, and prey per-area production *P_p_*, per-area death rate due to aggressiveness for prey and predators (*D_p_* and *D_π_*), the density of aggressive (*ρ_p_*_(_*_h_*_)_ and *ρ_π_*_(_*_h_*_)_) and nonaggressive (*ρ_p_*_(*d*)_ and *ρ_p_*_(*d*)_) prey and predators, and their per-area birth rate (*B*) are shown. (a) shows primary scaling laws, and (b) shows primary fluctuation scaling laws, according to which the fluctuation of the same variables (shown by *σ*) scales with *λ*. Parameter values: *η* = 0.05, *γ* = 0.5, *d* = 2, *q* = 0.5. An ecoevolutionary model with *n_g_* = 1, *r* = 0.4, *d_t_* = 0.05, and *ν* = 10*^−^*^3^ is considered.

The primary scaling relations have profound consequences. For any two generic variables, *v* and *w*, with the property *v* ∝ *λ^a_λ,v_^*, *w* ∝ *λ^a_λ,w_^*, we have *v* ∝ *w^a_λ,v_/a_λ,w_^*. This observation gives rise to a rich set of mean and fluctuation scaling laws. In Fig. 5, we decompose these scaling laws into four classes. The predator-prey scaling laws state that a predator variable scales with a prey variable. Some of these scaling laws are presented in Fig. 5(a) and include a recently discovered predator-prey power law according to which the density of predators as a function of the density of prey shows a power law, and prey production power law, according to which prey production increases with a sublinear power of prey density [12]. Furthermore, the model predicts prey and predator birth increases as a power of their density.

**Fig 5.**
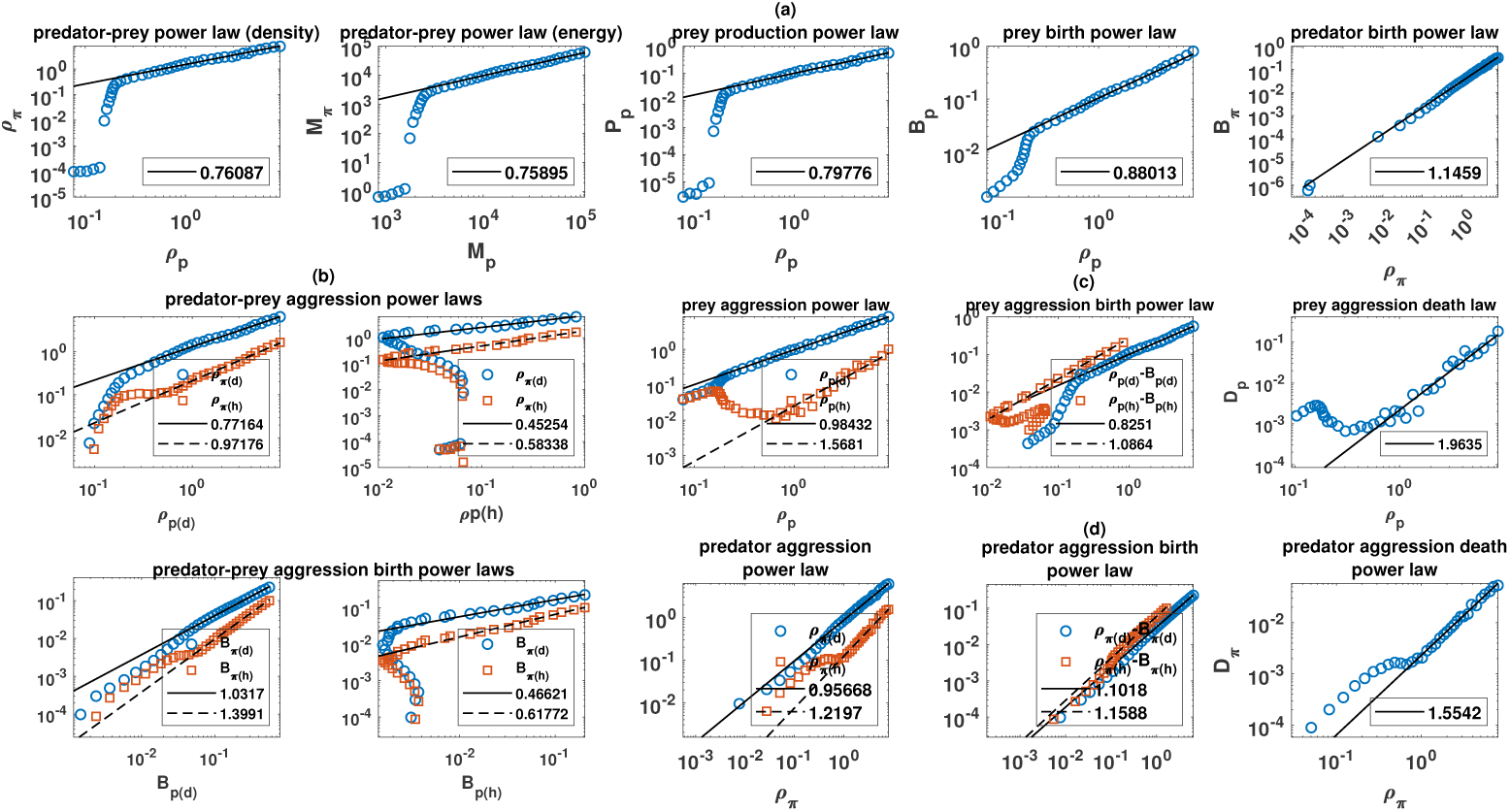
Eco-evolutionary scaling laws. (a): Predator-prey scaling laws include the scaling of predator density, *ρ_π_* and prey production, *P_p_* with prey density *ρ_p_*, and the scaling of the per-area birth rate of prey, *B_p_* and predators, *B_π_* with their density. (b): Predator-prey aggression scaling laws state that the density and birth rate of subpopulations of predators with different aggressiveness is a power law of the density and birth rate of prey with different aggressiveness. Aggressive predators have a higher exponent and grow faster than nonaggressive ones. (c): Prey aggression power laws state that the density of prey with different aggressiveness, *ρ_p_*(*h*) and *ρ_p_*(*d*) is a power law of total prey density *ρ_p_*, with a higher exponent for aggressive preys (left). Similarly, per-area birth rate of aggressive and nonaggressive prey versus their density, *ρ_p_*(*h*) − *B_p_*(*h*) and *ρ_p_*(*d*) − *B_p_*(*d*), is a power law (center). Furthermore, the per-area death rate due to aggressiveness increases superlinearly with prey density, which implies an increasing cost of aggressiveness in higher densities (right). (d): Predator aggression power laws state that the density of aggressive *ρ_π_*(*h*) and nonaggressive *ρ_π_*(*d*) predators is a power law of total predator density *ρ_π_*, with a higher exponent for aggressive predators (left). Similarly, aggressive and nonaggressive predator birth versus their density, *ρ_π_*(*h*) − *B_π_*(*h*) and *ρ_π_*(*d*) − *B_π_*(*d*), shows a power law with a higher exponent for aggressive predators (center). The per-area death rate of predators due to aggressiveness increases superlinearly with predator density (right). Parameter value: *η* = 0.05, *γ* = 0.5, *d* = 2, *q* = 0.5. An ecoevolutionary model with *n_g_* = 1, *r* = 0.4, *d_t_* = 0.05, and *ν* = 10*^−^*^3^ is considered.

Predator-prey aggression or competition scaling laws, presented in Fig. 5(b), relate the density or birth rate of subpopulations of prey and predators with different aggressiveness. According to these power laws, the density of aggressive or nonaggressive predators increases with the power of the density of aggressive and nonaggressive predators. The scaling exponent increases by increasing the aggressiveness of the predator and decreases by increasing the aggressiveness of the prey. This suggests that aggressive individuals grow faster. A point to which we will return shortly.

Finally, scaling laws exist within the same population. These scaling laws can be called aggression or competition power laws and are presented in Fig. 5(c) for prey and in Fig. 5(d) for predators. Aggression scaling laws hold for subpopulations and state that aggressive and nonaggressive individuals increase according to a power law of the total density of the population (top right panel in Fig. 5(c) for prey and Fig. 5(d) for predators). Furthermore, per-area birth rate of aggressive and nonaggressive individuals is a power law of their density with a higher exponent for higher aggressiveness (top middle panel in Fig. 5(c) for prey and Fig. 5(d) for predators). Finally, the per-area death rate due to aggressiveness increases with a super linear power law with density (top left panel in Fig. 5(c) for prey and Fig. 5(d) for predators).

In addition to the scaling of means, the primary scaling relations imply scaling of the temporal fluctuation of a variable with the mean or the temporal fluctuation of another variable (See SI. 8 and SI. 9). These scaling relations extend the famous Taylor’s power law, according to which the temporal or spatial fluctuation of populations is a power law of their mean, to what can be called heterogeneous fluctuation-scaling laws, according to which fluctuation and mean of different variables pertaining to the same population, or even different populations or subpopulations within a population scale with each other. The scaling exponent of the heterogeneous fluctuation scaling power laws is often close to 1, which indicates strong dependencies and correlations in the system [24] in the non-equilibrium self-organized critical regime.

### The evolutionary dynamics: Out of equilibrium eco-evolutionary dynamics gives rise to heterogeneous social environment and consistent personalities

In Fig. 6, we begin to study the evolutionary dynamics of the system by plotting the fraction of prey and predators with aggressive and nonaggressive strategies in a non-viscous (Fig. 6(a)) and viscous population (Fig. 6(b)). Here, we have taken the population viscosity of prey and predators to be the same, which does not need to be always the case (see below). Comparison between viscous and non-viscous populations shows that population viscosity significantly decreases aggressiveness. Furthermore, while in non-viscous populations, predators evolve a higher aggressiveness, preys evolve to be more aggressive than predators in a viscous population. Both phenomena can be understood in terms of the scarcity of resources. In a non-viscous population, resources for predators are subject to higher uncertainty, thus more valuable and giving rise to higher resource competition. However, population viscosity leads to a similar resource competition in prey, which weakens predator-prey differences in evolving aggressive strategies. At the same time, population viscosity leads to a higher chance of interaction with close relatives, which suppresses aggressive strategies. Consequently, despite higher resource competition in a viscous population, a lower aggressiveness in both prey and predators is observed. This is reminiscent of the beneficial effect of limited motility on the evolution of cooperative behavior [43, 44].

**Fig 6.**
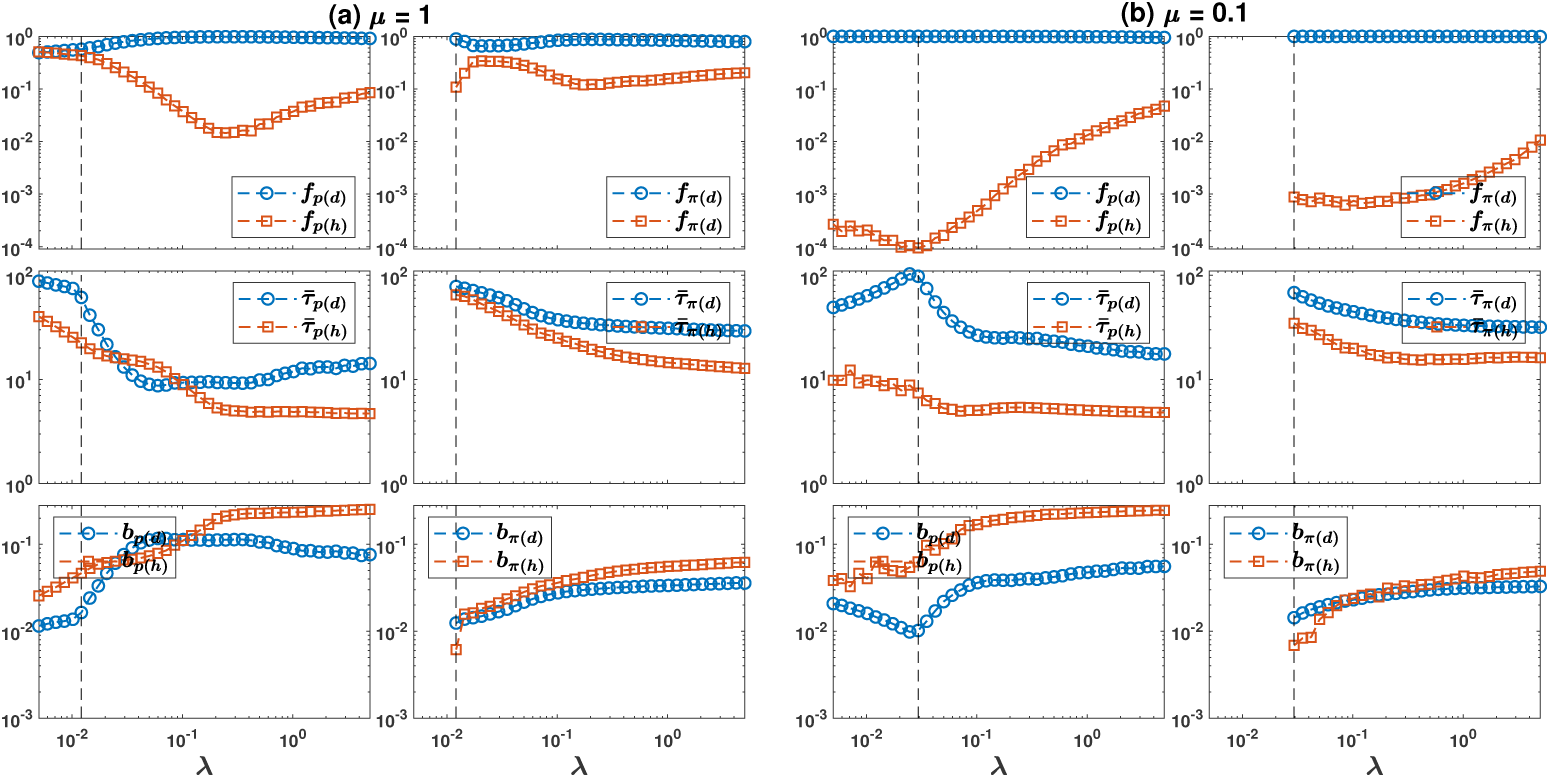
The evolution of aggressiveness and life-history trade-offs. From top row to bottom, the fraction, *f*, mean lifespan, *τ̄*, and the per-capita birth rate, *b*, of nonaggressive and aggressive prey (left) and predators (right) are plotted. In (a), a nonviscous population with movement probability *µ_p_* = *µ_π_* = *µ* = 1, and in (b), a viscous population with movement probability *µ_p_* = *µ_π_* = *µ* = 0.1 is considered. By increasing the density, aggressiveness decreases in low densities (equilibrium regime) and increases in high density (non-equilibrium regime). Ecological dynamics give rise to the evolution of life history trade-offs according to which aggressive individuals live shorter lives and reproduce more than nonaggressive ones. Population viscosity hinders the evolution of aggressiveness. Parameter value: *η* = 0.05, *γ* = 0.5, *d* = 2, *q* = 0.5. An ecoevolutionary model with *n_g_* = 1, *r* = 0.4, *d_t_* = 0.05, and *ν* = 10*^−^*^3^ is considered.

An interesting prediction of the model is the evolution of life history differences between aggressive and nonaggressive strategies. In both a viscous and non-viscous population, aggressive individuals have a shorter average lifespan and higher per-capita reproduction (Fig. 6 (middle row)). This shows that aggressive and nonaggressive individuals evolve different life histories. Nonaggressive individuals avoid the risk of death in confrontation at the expense of losing resources. This leads to a higher lifespan. Aggressive individuals obtain higher resources in confrontations and, thus, enjoy a higher per-capita birth at the expense of a higher death rate. Importantly, life history trade-offs are weaker for preys. As we will shortly see, this point has important consequences.

One might wonder if such life history trade-offs can lead to the evolution of consistent personalities. This can be the case due to the fact that, since aggressive individuals develop life-history traits adapted to aggressiveness, i.e., higher reproduction and lower lifespan, acquiring traits that further decreases their lifespan is not as detrimental for them as it is for nonaggressive individuals who invest in having a long lifespan [40, 41]. To examine this idea, we consider the model with *n_g_* = 2, i.e., allow individuals to play two games, where their strategies in the two games is a priori independent. The correlation between the strategies of the individuals in the two games, which grasps the repeatability of individuals’ behavior and is core to animal personalities [40, 41, 54, 55], is plotted in Fig. 7(a). Interestingly, the correlation between the strategies of the individuals in the two games becomes negative in the equilibrium regime and positive in the self-organized critical regime. A negative correlation implies that individuals who are aggressive in one game are more likely to be nonaggressive in another game. This allows individuals to anti-coordinate in a game equilibrium where they defer to others in one game for others to defer in another game. As shown in the Methods, such a reciprocal strategy can be a Nash equilibrium of the combination of the two games when the expected lifespan is long enough, and thus, death is highly costly, which is what holds in the small density regimes (Fig. 6 middle row).

**Fig 7.**
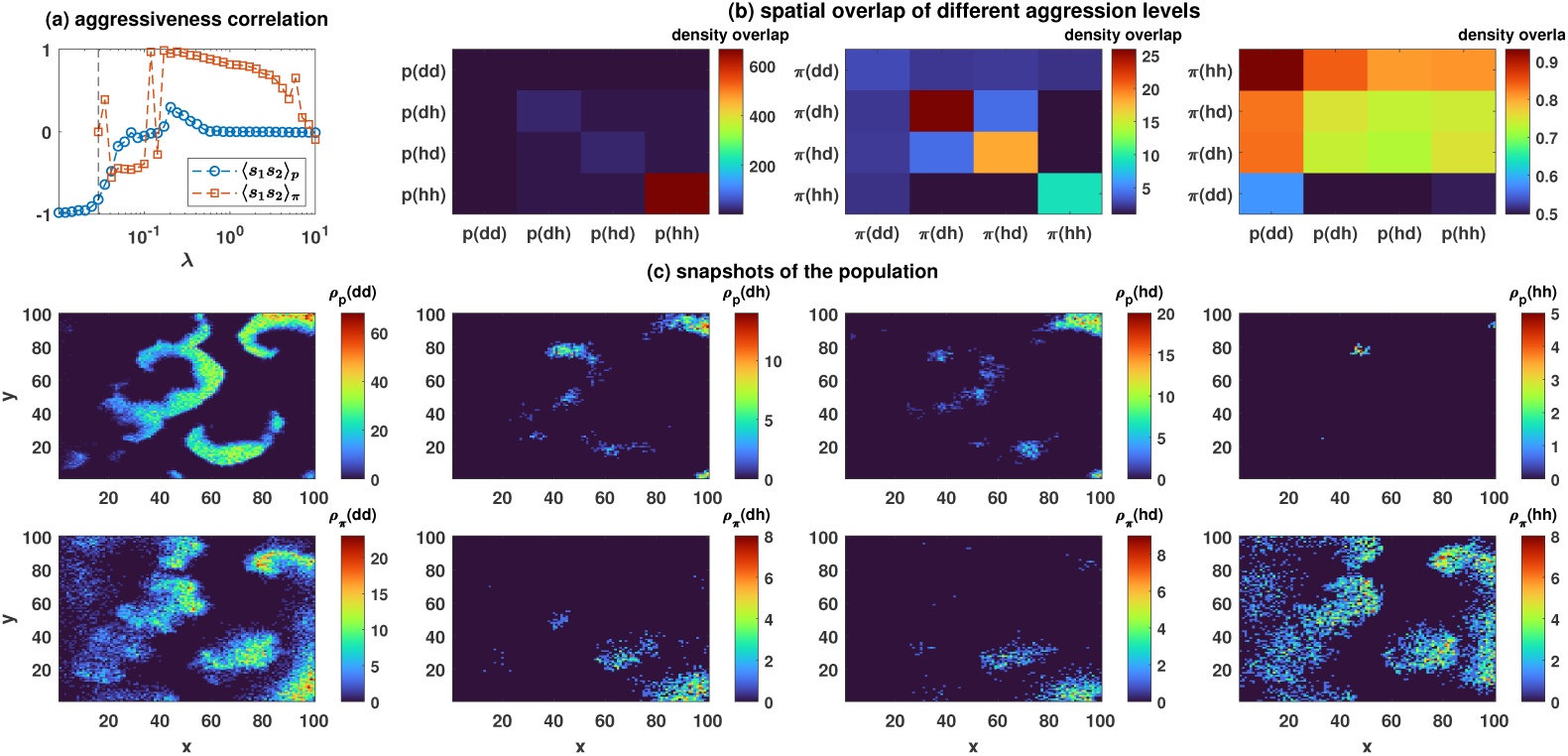
The evolution of consistent personalities and heterogeneous social environment in the out-of-equilibrium eco-evolutionary dynamics. (a): The correlation between the strategies of the individuals in the two games, separately for preys and predators, is plotted. (b): normalized spatial overlap between the densities of the individuals in the two games for preys (left), predators (center), and between preys and predators (right) in the non-equilibrium regime is color plotted. (c): Snapshots of the spatial distribution of prey (top) and predators (bottom) with different aggressiveness in the non-equilibrium regime. In small densities with equilibrium dynamics, individuals anti-coordinate in a Nash equilibrium where they evolve to be aggressive in one game and defer in another game, which leads to a negative correlation between strategies in the two games. In contrast, in large densities with non-equilibrium dynamics, a positive correlation between the strategies of the individuals in the two games evolves, which shows the evolution of consistent personalities (a). Spatial distribution of the individuals reveals that while heterogeneous strategies, *dh* and *hd*, tend to occupy the same spatial locations where they benefit from anti-coordination, nonagressive (*dd*) and highly agressive (*hh*) individuals tend to co-occure in space. This creates a heterogeneous social environment where individuals with different personalities and life histories coexist. Aggressive individuals invest in maximizing reproduction, and nonaggressive ones invest in maximizing lifespan. Comparison of prey and predators show that heterogeneous social environments and consistent personalities are much stronger in predators. This is due to the fact that while for predators, aggressiveness increases the density overlap between prey and predators, for prey, the density overlap decreases with aggressiveness. This decreases the risk of death due to predation for aggressive prey and increases their lifespan, which weakens prey life history trade-offs and personality consistency. Parameter value: *η* = 0.05, *γ* = 0.5, *d* = 2, *q* = 0.5. An eco-evolutionary model with *n_g_* = 1, *r* = 0.4, *d_t_* = 0.05, and *ν* = 10*^−^*^3^ is considered.

On the other hand, as lifespan decreases (high *λ*), the cost of death decreases, and a simple argument suggests *hh* becomes the Nash equilibrium of the game (Methods). However, this contrasts with what is observed in the simulation. Rather than settling in a homogeneous Nash equilibrium, we observe a heterogeneous population of individuals with consistent personality differences. The deviation from the Nash equilibrium results from the heterogeneity in the social environment that individuals experience, which, in turn, originates from the non-equilibrium self-organized critical dynamics of the system. To see this, in Fig. 7(c), we plot snapshots of the population for prey (top row) and predators (bottom row) with different strategies when *n_g_* = 2. *hd* and *dh* individuals are unlikely to co-occur with *hh* strategies and are more likely to coexist in the same spatial domains, where they benefit from anti-coordination to avoid a high risk of death. On the other hand, *hh* strategies are more likely to appear in the same domains as *dd* individuals, where they can exploit the latter. However, too high a density of *hh* individuals leads to a high cost of aggressiveness and keeps the population of aggressive individuals in check. The fact that individuals with different strategies experience contrasting social environments is also apparent in Fig. 7, where the overlap between the density field of individuals with different strategies is plotted. We note that these results hold for other values of *λ*, and the density overlaps are also among the variables which show scaling with *λ*.

An interesting observation is that aggressive preys have a lower overlap with predators and, thus, a lower risk of being predated. In contrast, aggressive predators have a higher overlap with prey and, thus, a higher chance of catching prey. The reason is that aggressive individuals tend to repel each other and are less likely to live in dense packs. While living in dense groups endows a shelter to prey, it also increases the chance of getting captured due to the fact that predators grow faster in prey-rich regions and effectively chase nonaggressive preys (see the Supplementary Videos). This leads to a higher overlap of predators and nonaggressive prey. On the other hand, for predators, living in diluted groups leads to a higher chance of catching prey due to lower competition and, thus, faster growth in regions where prey are abundant, which leads to a higher overlap with the prey density field. This argument has three important consequences. First, it shows that both prey and predator aggressiveness can have indirect and surprising benefits, which favors its evolution. Second, it suggests that competition between prey can surprisingly benefit prey growth. We will shortly return to this point. Lastly, a lower predation risk of aggressive prey compensates for a higher risk of death in aggressive encounters and increases aggressive prey mean lifespan. This weakens preys’ life history trade-offs, which in turn leads to a weaker personality consistency in preys.

### Factors affecting the predator-prey growth

Scaling relations are universal and robust features of ecological and eco-evolutionary dynamics and hold irrespective of whether evolution takes place or not and are surprisingly robust under different modifications of the model. To argue for this point, in addition to analyzing different parameter values in the Supplementary Information (SI. 9), we consider modifications of the model with or without evolution, with different motility, or where offspring size is smaller than that of the parent, and in all the cases the same self-organized critical dynamics with scaling laws is observed. While scaling relations are universal, the value of the scaling exponents depends on the parameter values or modifications of the model. Here, we study some of the factors affecting one of the scaling exponents, the predator-prey power law, more explicitly, given that this power law has been recently observed in ecosystems and seems to be a sublinear power law [12].

To see how predator competition affects predator-prey power law, in Fig. 8(a), from top to bottom, we plot the density of predators as a function of the density of prey, predator total internal energy as a function of prey total internal energy, and prey production as a function of prey density, for different values of *n_g_*. The predator-prey power law exponents decrease by increasing the extent of competition. This pattern is also valid when only predators compete or when the extent of competition is controlled by varying the reward, *r* (see the Supplementary Information, SI. 8 and SI.9), and shows that predator competition decreases the predator-prey power law exponent and contribute to a bottom-heavy food web. As shown in the Supplementary Information (SI. 8), surprisingly, prey competition can also favor prey growth and decreases the predator-prey power law exponent. This results from a more scattered spatial distribution of aggressive prey. However, the beneficial effect of prey competition is compensated by its direct negative effect on prey growth due to a higher death rate in aggressive encounters and can only lead to a weak benefit for prey growth.

**Fig 8.**
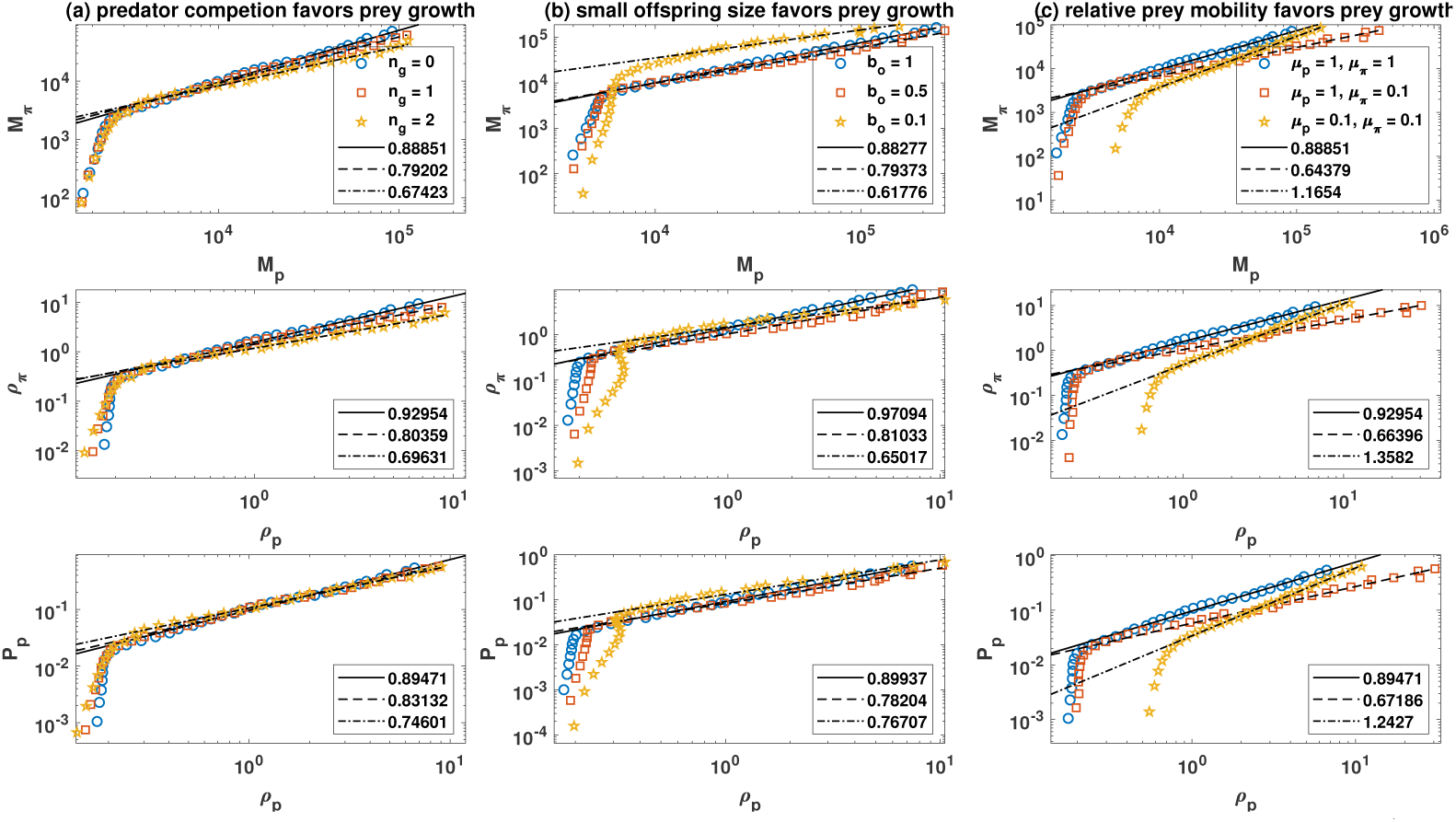
Determinants of predator-prey power law exponent. Predator total energy (top row) as a function of prey total energy, and predator density (middle row) and prey production defined as the number of prey caught by predators (bottom row), as a function of prey density, for different numbers of games, *n_g_*, (a), different offspring size, *b_o_*, (b), and different motility of prey, *µ_p_* and predator, *µ_π_*, (c), are plotted. (a): Increasing the number of games increases the extent of competition and leads to smaller predator-prey power laws, which shows predator competition favors prey growth. (b): In a model where individuals transfer an amount *b_o_* of their energy to their offspring, by decreasing the offspring size, the predator-prey power law exponent decreases, which shows small offspring size favors prey growth. (c): High relative prey motility (*µ_p_* = 1, *µ_π_* = 0.1) favors prey growth and decreases the predator-prey law exponent. Limited motility of both prey and predator (*µ_p_* = 0.1, *µ_π_* = 0.1), by limiting the escaping capacity of prey, favors predator growth and leads to a superlinear scaling. Parameter value: *η* = 0.05, *γ* = 0.5, *d* = 2, *q* = 0.5. In (b) and (c) an ecological model (*n_g_* = 0) is considered and in (a), *r* = 0.4, *d_t_* = 0.05, and *ν* = 10*^−^*^3^.

We examine the effect of varying the offspring size by considering a modification of the model where, instead of the resource of the parent to be divided in half upon reproduction, a constant amount *b_o_* is transferred to the offspring, in Fig. 8(b). This model is studied in depth in the Supplementary Information and shows similar phenomenology to the model considered so far. As here the reproduction threshold is taken equal to *d* = 2, *b_o_* = 1 requires, on average, the parent’s resources to be halved between the parent and offspring, and is similar to the model with mitosis. This case yields a similar exponent to that observed in the model with mitosis. However, decreasing the offspring size leads to a smaller predator-prey power law exponent.

Comparison with the model with mitosis in the Supplementary Information (SI. 8), shows that with smaller offspring size prey density grows faster by increasing the resources, while predator birth and density grow slower. This is due to the fact that a large number of predator kills are prey offspring of small sizes, which reduces the flow of resources to predators and leads to slower growth of predators. Moreover, the exponents of the spatial distribution (*ν*) of prey decreases, and that of predators increases with smaller offspring size. This implies with smaller offspring sizes, large number of prey on the same lattice site are observed more frequently, and large number of predators are observed less frequently. Furthermore, the exponent of internal energy of both prey and predators with *λ* significantly decreases with small offspring size. This result shows that with smaller offspring sizes, biomass (of both prey and predators) shows slower growth with increasing resources.

In Fig. 8(c), we study how motility affects the predator-prey power law. Small relative predator motility (*µ_p_* = 1 and *µ_π_* = 0.1) leads to a smaller predator-prey power law exponent. In contrast, high predator relative motility can lead to the instability of the system in high resources. When both prey and predator populations are viscous, the predator-prey power law can even become a superlinear power law. A sublinear predator-prey power law suggests a bottom-heavy food web pyramid and bottom-up control of the food web [12]. Examination of the direction of control based on the information flow [56] from predator to prey birth rate in the Supplementary Information (SI. 7) shows this is indeed the case and establishes a link between prey and predators’ life histories, the direction of control in the food web [57–60], and the sublinear or superlinearity of the predator-prey scaling. A bottom-heavy food web with sublinear predator and prey production growth occurs when preys control the food web. In this case, a prey-favoring life history density dependence occurs, according to which the mean lifespan of prey increases by increasing the density, and their per-capita birth decreases. This, in turn, results from the fact that due to higher density fluctuations in higher densities, predation becomes more difficult [25]. On the other hand, limited prey motility limits the prey’s ability to escape predators, and predators can consume all the prey when they arrive in a neighborhood. While it is intuitively expected that this leads to the predator control of the food web, the model shows, combined with the scale-invariance spatial distribution of predators and prey, low motility leads to a superlinear scaling of predator density with prey density. In this case, a predator favoring life history density dependence occurs, according to which by increasing the density, prey per-capita production increases and their mean lifespan decreases.

In the Supplementary Information (SI. 9), we show that a lower predator conversion efficiency and small body size also lead to slower predator growth with increasing density and smaller predator-prey power law exponents. Finally, the model has important predictions for life histories and their density dependences (SI. 7 and SI. 9). For instance, the model shows that small animals reproduce more and live shorter lives due to their susceptibility to starvation. Importantly, small body size, competition, and smaller offspring size lead to faster life history patterns in higher densities (shorter lifespan and higher per-capita birth rate) in both prey and predators. On the other hand, lower predator conversion efficiency and higher motility lead to slower life histories in higher densities for prey. For predators, a lower conversion efficiency leads to faster life history, and motility has a low effect on predators’ life history. These findings suggest scaling relations in the life histories of individuals may be at work and relate to the food web control and population scaling relations.

## Discussion

As we have seen, a simple model of ecological interactions based on the flow of resources in ecosystems provides a natural framework for evolutionary phenomena. This approach suggests that flow and dissipation of energy in ecosystems can provide a unifying framework for understanding ecological and evolutionary dynamics in which ecosystems can be regarded as self-organizing complex systems [32–35], and ushers the conception of an eco-evolutionary approach which can grasp large-scale eco-evolutionary patterns [30, 31]. The model suggests trophic interactions lead to the self-organization of ecosystems into a critical state manifested by power-law distributed avalanches of activity and scale-invariant frequency-size distributions. Scaling laws are an important consequence of this scale invariance: the model suggests the recently discovered predator-prey power law is just the tip of an iceberg of scale-invariant ecosystems, and these power laws can be extended to a wide range of scaling laws. Furthermore, the model provides theoretical evidence for a new class of scaling laws, aggression scaling laws which hold in and between populations. These theoretical predictions call for new empirical research to uncover the consequences of scale-invariance of eco-evolutionary dynamics. Furthermore, while some of the predator-prey power laws reported here have been recently discovered in large ecosystems [12], the model suggests predator-prey power laws can be universally at work, potentially even with a superlinear exponent, in simple trophic communities such as microorganisms, which controlled lab experiments can test.

The question of whether food webs show top-down [57] or bottom-up control [58] has been subject to much research [58–60]. Recent work recognizes that both bottom-up and top-down forces can be at work in real food webs, and their relative strength determines the food web control [58–60]. For instance, past work has argued that a prey-dependent functional response leads to a top-down food web where increasing resources increase predator density but not prey [61, 62]. The model predicts this can happen in small densities where predation can be approximated by a linear response to prey density [25]. However, it is argued that the factors which limit predator capacity to consume prey, which can require a ratio-dependent functional response, can change the nature of control in the food webs [58, 59, 63]. The model shows this is the case and identifies several forces that can strengthen bottom-up or top-down control. For instance, consistently with past work, the model shows predator competition favors bottom-up control [63, 64]. The fact that low relative predator motility and low conversion efficiency favor bottom-up control also seems to conform with the idea that factors impeding predator’s efficiency in exploiting prey contribute to bottom-up control [59]. Besides, the model predicts small offspring size and small body size can contribute to bottom-up control.

Importantly, the model establishes a link between food web control and life history patterns and scaling in food webs by showing that the same factors contributing to a bottom-up control can lead to smaller predator-prey power law exponent and slower prey life history patterns in higher densities. Given that the density dependence of life history patterns has been subject to much theoretical and empirical interest [45–47], further investigation of the link between life history patterns, food web control, scaling laws, and self-organized criticality of the eco-evolutionary dynamics can lead to an important cross-fertilization of the theoretical and empirical research in diverse ecological disciplines.

The eco-evolutionary approaches also shed light on the evolution of life history traits, trade-offs, and consistent personalities [40, 41, 55, 65]. The model shows predators evolve to be more aggressive than prey and show a higher personality consistency. Furthermore, aggressiveness can have indirect benefits by increasing predation for predators and decreasing the predation risk for prey, which favors its evolution. Life history trade-offs can emerge naturally as a result of eco-evolutionary dynamics and are a necessary but not sufficient condition for the evolution of a consistent personality. Rather, deviation from a Nash equilibrium and the evolution of consistent personalities crucially depends on the heterogeneous social environment brought about by the out-of-equilibrium self-organized critical dynamics. Finally, the model suggests population viscosity hinders the evolution of aggressiveness and personality consistency by deriving the evolution of homogeneous nonaggressive populations due to the beneficial effect of population viscosity on the evolution of altruistic behavior [43, 44].

## Methods

### The simulations

At each time step of the simulation, the lattice sites are updated sequentially. At first, preys consume the resources available on the lattice site. For this purpose, all the resources on a site are equally divided among the prey. We assume a prey can acquire up to *d* amount of resource per time step. *d* is the energy of the individual upon reproduction and can be understood as the adult size. This assumption is meant to make the model more realistic by setting a bound on the amount of resources an individual can acquire per time step. However, the overall phenomenology of the model is similar in the absence of this assumption.

In the next step, all the predators are given a chance to attempt predation. This is continued until either all the predators have attempted capturing a prey, or no more prey on the site exists. If a predator successfully captures a prey, it gains *γϵ_p_* amount of resource. Here, *ϵ_p_* is the internal energy of the captured prey. Then individuals are paired at random to play *n_g_* Hawk-Dove games. In this game, when two dove individuals meet, nothing happens. When a hawk meets a dove, an amount of resource *r* is transferred from the dove to the hawk. When two hawks meet, the net transfer of the resource is zero, and each individual dies with a probability *d_t_*. If *n_g_ >* 1, individuals are paired separately for each of their games. If an odd number of individuals live on a site, one of the individuals (determined by chance) is left with no opponent and does not play the game.

After playing the game, individuals reproduce. In this stage, all those whose internal resource is higher than *d* produce an offspring. In the model with mitosis, the parent’s resource is divided equally among the parent and the offspring. In the model with variable offspring size, the parent always transfers *b_o_* of its resource to the offspring.

After reproduction, individuals move. In this stage, each individual moves to one of the randomly chosen neighboring sites on the lattice with probability *µ_p_* for prey and *µ_π_* for predators. Unless otherwise stated, we take *µ_p_* = *µ_π_* = 1. In the last stage, individuals pay their metabolic cost, *η*, and all those whose internal energy is equal to or smaller than the subsistence threshold Δ are removed from the population. Unless otherwise stated, we set Δ = 0 to reduce one of the parameters.

Finally, the amount of resources available on a site increases by an amount *λ*(***r***). We have used a linear distribution of resources. That is, the resource regeneration rate on a site ***r*** = (*x, y*) is given by *λ*(*r*) = 2*λx/L*, where *x* is the *x*-component of the position, ***r***. The factor of 2 ensures the average resources regeneration rate in the system is equal to *λ*. In the Supplementary Information (SI. 3), we also consider a homogeneous distribution of resources where the resource regeneration rate on all the lattice sites is the same and equals *λ*. Resource distribution can affect the stability of the system.

However, different ways of implementing resource heterogeneity (such as considering a patchy environment [25]) can stabilize the system, and the results are independent of the details of the distribution of the resources.

For the spatial structure, we have considered a first nearest neighbor square lattice with von Neumann connectivity and fixed boundaries with linear size *L*. In the simulations performed in the main text, we set *L* = 100, and in the Supplementary Information, *L* is varied from *L* = 100 up to *L* = 200. We have performed simulations for different initial numbers of prey and predators with randomly distributed hawk and dove strategies. The fate of simulations in the purely ecological model is independent of the initial conditions. This results from the ergodicity of the dynamics. Unless otherwise stated, the simulations start with 100 prey and 10 predators. In the eco-evolutionary model with zero or too small mutation rates, the fraction of aggressive and nonaggressive prey and predators can depend on the initial conditions. As shown in the Supplementary Information (SI. 5), this results from the fact that aggressive preys go to extinction for too low a resource regeneration rate and small system size.

As mentioned before, when *n_g_* = 0, the model reduces to a purely ecological model of trophic interactions. This model has the elegant property of giving rise to a natural framework for evolution by allowing individuals to have inheritable traits. This property is exploited in the eco-evolutionary model where individuals play *n_g_ >* 0 games. In this way, by changing *n_g_* from zero, one can turn on the evolution in the model. As shown in the supplementary information, all the results are valid for both the ecological and evolutionary versions of the model.

**Table 1.** The game payoffs.

|  | dd | dh | hd | hh |
| --- | --- | --- | --- | --- |
| dd | 0,0 | $-r, r$ | $-r, r$ | $-2r, -2r$ |
| dh | $r, -r$ | $-c_d, -c_d$ | 0,0 | $-r - c_d, r - c'_d$ |
| hd | $r, -r$ | 0,0 | $-c_d, -c_d$ | $-r - c_d, r - c'_d$ |
| hh | $2r, -2r$ | $r - c'_d, -r - c_d$ | $r - c'_d, -r - c_d$ | $-2c'_d, -2c'_d$ |

### Nash equilibrium of the game with ***n_g_* = 2**

In the modified version of the Hawk-Dove game used here, agents may receive a numeric value in terms of resources or face a risk of death from the game they play. To write down a payoff matrix for the game, one needs to estimate a numerical value for the risk of death in terms of its equivalent resource loss. Without loss of generality, assume the cost equivalent of a risk of death for *hh* strategy is equal to *c^′^* and for *dh* and *hd* strategies is *c_d_*. We will shortly comment on how to relate these parameters with model parameters. The payoff matrix of the game in terms of the cost equivalent of the risk of death for *n_g_* = 2 is presented in Table. 1.

When *r < c_d_^′^*, the game has two pure strategy Nash equilibrium, (*hd, dh*) and (*dh, hd*). This equilibrium is similar to the anti-coordination equilibrium in the simple Hawk-Dove game. On the other hand, for *c^′^_d_ < r* and *c_d_^′^ <* (*r* + *c_d_*)*/*2, (*hh, hh*) becomes the Nash equilibrium. The second inequality can be written as *c_d_^′^ < r* + 2(*c_d_* − *c_d_^′^*), and assuming (*c_d_* − *c_d_^′^*) *>* 0, which as explained shortly empirically holds, is satisfied when c*_d_^′^ < r*. This simplfies the behavior of the game: as the cost of death for *hh* decreases below the reward, *r*, *hh* becomes the Nash equilibrium of the game.

Now we need to relate the cost equivalent of death to the risk of death. To do so, note that the cost of death is an inverse function of the expected lifespan. As shown in the supplementary information (SI. 5), the expected lifespan of *dh* (which is the same as the expected lifespan of *hd*) is slightly smaller than the expected lifespan of *hh*. This shows that the cost of death for *dh* is higher than the cost of death for *hh*, *c_d_^′^ < c_d_*, and justifies our assumption above. Furthermore, the expected lifespan monotonically decreases by increasing *λ*. Thus, both *c_d_^′^* and *c_d_* decrease by increasing *λ*. Thus, for small *λ*, the anti-coordination equilibrium is the Nash equilibrium of the game, and as *λ* increases, *hh* becomes the Nash equilibrium of the game.

Simulation results show that for sufficiently small *λ*, where the lifespan of prey is too high, the anti-coordination equilibrium indeed occurs with high motility. However, simulations show that the Nash equilibrium *hh* does not occur for large *λ*. Rather, a heterogeneous population where all the strategies coexist is observed. The deviation from the Nash equilibrium results from the non-equilibrium dynamics, which introduces density fluctuations in the system. This increases the assortativity of the interactions and favors nonaggressive strategies, as they are more likely to interact with fellow nonaggressive individuals (similarly, limited motility by giving rise to assortative interactions leads to a deviation from the Nash equilibrium). This argument shows that the non-equilibrium dynamic is essential for a deviation from the Nash equilibrium and the evolution of consistent personalities.

### Relating the spatio-temporal scale invariant distribution of mass with scaling laws: A mass-scaling theory

Extending the theory developed in [25], here we show that the power law distribution of a generic quantity, *x*, together with the scaling of its cut-off with resource regeneration rate leads to the scaling of *x* with *λ*. Let *f* (*n_x_, λ*) be the probability of observing the value *n_x_* on a lattice site for a fixed *λ*. We have for the average value of *x* on a lattice site, 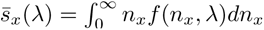 (when *x* is a discrete variable, we have 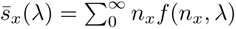, which can be approximated by an integral using Euler-Maclaurin theorem). Taking the lower cut-off equal to zero, and using the simple scaling form, 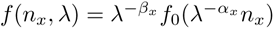, we have 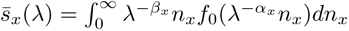. Assuming *f*_0_ exhibits a sharp cut-off, the infinity in the upper bound of integral can be replaced by the cut-off, 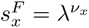 and assuming *f*_0_(*n_x_*) has a power law form, 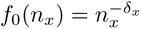, we have:

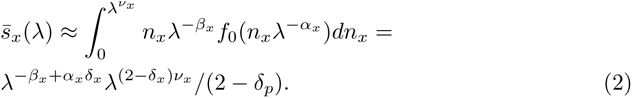

Thus, the theory predicts *s̄_x_* obeys scaling with *λ* with an exponent, 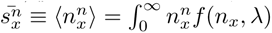. A similar calculation for the *n*th moment of *x*, 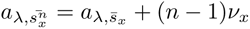, yields, 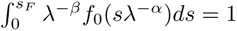. We note that the normalization condition on *s̄_x_* predicts a Taylor power law with strong density dependence: neglecting the mass of the probability density function above the upper cut-off and taking the lower cut-off equal to zero, we have 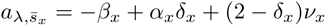, which gives −*β_x_* + *α_x_δ_x_* = −(1 − *δ_x_*)*ν_x_*. This implies 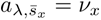, which predicts a spatial fluctuation scaling law for *x* with an exponent of one, 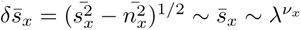 that is a highly aggregated dynamics observed in the non-equilibrium regime.

Our estimates for the exponents for primary resources, prey, and predators are as follows: *δ_r_* = 0.35, *δ_p_* = 0.5, and *δ_π_* = 0.85. *ν_r_* = 1.23, *ν_p_* = 0.55, *ν_π_* = 0.52, *α_r_* = 1, *β_r_* = 1, *α_p_* = 0.75, *β_p_* = 0.65, *α_π_* = 0.75, *β_π_* = 0.8. These estimates predict the mass scaling exponent of prey density: *a_λ,ρ_p__* = 0.55, and for predator density, *a_λ,ρ_p__* = 0.41, which predicts a predator-prey density scaling exponent of *a_ρ_p_,ρ_π__* = 0.745. The estimates based on a linear fit in a log-log plot for these exponents are: *a_λ,ρ_p__* = 0.77, *a_λ,ρ_p__* = 0.60, and *a_λ,ρ_p__* = 0.81. Therefore, the scaling theory predicts the primary scaling exponents with an error of up to 30 percent and the predator-prey power law exponent with an error of less than 10 percent.

Similarly, as shown in the Supplementary Information (SI. 4), the distribution of prey and predators with dove and hawk strategies obeys a power law. The exponents of the power law distribution are estimated as follows: *δ_p_*_(*d*)_ = 0.52, *δ_p_*_(_*_h_*_)_ = 0.99, *δ_π_*_(*d*)_ = 0.87, *δ_π_*_(_*_h_*_)_ = 1.05. Furthermore, the maximum number of individuals of a subpopulation observed on a lattice site obey a power law with the following exponents: *ν_p_*_(*d*)_ = 0.57, *ν_p_*_(_*_h_*_)_ = 0.76, *ν_π_*_(*d*)_ = 0.5, *ν_π_*_(_*_h_*_)_ = 0.44. The data collapse exponents are estimated as follows: *α_p_*_(*d*)_ = 0.75, *β_p_*_(*d*)_ = 0.65, *α_p_*_(_*_h_*_)_ = 0.95, *β_p_*_(_*_h_*_)_ = 0.65, *α_π_*_(_*_d_*_))_ = 0.75, *β_π_*_(*d*)_ = 0.8, *α_π_*_(_*_d_*_))_ = 0.55, *β_π_*_(_*_h_*_)_ = 0.55. These estimates predict for the primary aggression scaling exponents: *a_λ,ρ_p(*d*)__* = 0.58, *a_λ,ρ_p(*h*)__* = 1.05, *a_λ,ρ_π__*_(*d*)_ = 0.41, *a_λ,ρ_π__*_(_*_h_*_))_ = 0.45. These estimates give for the secondary aggression exponents: *a_ρ_p_,ρ_p(*d*)__* = 1.05, *a_ρ_p_,ρ_p(*h*)__* = 1.90, *a_ρ_p_,ρ_p(*d*)__* = 1, *a_ρ_p_,ρ_p(*d*)__* = 1.09.

The estimates of primary aggression exponents based on a linear fit in a log-log plot are *a_λ,ρ_p(*d*)__* = 0.75, *a_λ,ρ_p(*h*)__* = 1.2, *a_λ,ρ_π__*_(*d*)_ = 0.58, *a_λ,ρ_π__*_(_*_h_*_)_ = 0.73. Thus, the errors are, respectively, 17%, 13%, 30%, and 38%. For the secondary exponents, a linear fit in a log-log plot gives *a_ρ_p_,ρ_p(*d*)__* = 0.98, *a_ρ_p_,ρ_p(*h*)__* = 1.56, *a_ρ_p_,ρ_p(*d*)__* = 0.95, *a_ρ_p_,ρ_p(*d*)__* = 1.21. Thus, the prediction error of the mass scaling theory for these exponents is, respectively, 7%, 21%, 5%, and 9%. In this case, again, the mass scaling theory provides a better estimate of the secondary exponents compared to its estimates of primary exponents. This can be attributed to the fact that in writing eq. 2, we used the approximations that the mass of the integral above the upper cut-off and below the lower cut-off can be neglected. While this can introduce a large error in the determination of the primary exponents, these errors partially cancel each other when two primary exponents are divided to derive a secondary exponent.

These errors can be attributed to the approximations involved in the theory (taking the lower cut-off equal to zero and neglecting the mass of the integral above the upper cut-off), as well as the errors involved in estimating the exponents based on a linear fit in a log-log plot. Another source of error relies on a reliable determination of the data collapse exponents, *α* and *β*. As we have relied on a visual examination of the data collapse, we expect our estimates of these exponents to be subject to a higher error compared to other exponents.

### The direction of information flow and food web control

We use the direction of information flow [56] between the time series of the birth rates of prey and predators to examine the food web control (see Supplementary Information, SI.7). Information flow from the birth rate of prey to predators can be defined as 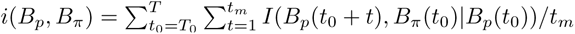. Here, *I* stands for conditional mutual information, and *B_p_*(*t*) and *B_π_*(*t*) are the time series of the per-area birth rate of, respectively, preys and predators at time *t*. The direction of information flow can be defined as the normalized difference of the information flow from the density of predators to prey and vice versa:

*D* = [*i*(*B_π_, B_p_*) − *i*(*B_p_, B_π_*)]*/*[*i*(*B_π_, B_p_*) + *i*(*B_p_, B_π_*)].

## Availability of Data and Materials

All data generated or analysed during this study are included in this published article and its supplementary information files.

## Acknowledgement

The author acknowledges funding form the Deutsche Forschungsgemeinschaft (German Research Foundation) under Germany’s Excellence Strategy-EXC 2117-422037984 during parts of this research.

## Notes

### Competing Interest Statement

The authors have declared no competing interest.

